# Induction of Dopaminergic Neurons for Neuronal Subtype-Specific Modeling of Psychiatric Disease Risk

**DOI:** 10.1101/2021.04.01.438094

**Authors:** Samuel K. Powell, Callan O’Shea, Kayla Townsley, Iya Prytkova, Kristina Dobrindt, Rahat Elahi, Marina Iskhakova, Tova Lambert, Aditi Valada, Will Liao, Seok-Man Ho, Paul A. Slesinger, Laura M. Huckins, Schahram Akbarian, Kristen J. Brennand

## Abstract

Dopaminergic neurons are critical to movement, mood, addiction, and stress. Current techniques for generating dopaminergic neurons from human induced pluripotent stem cells (hiPSCs) yield heterogenous cell populations with variable purity and inconsistent reproducibility between donors, hiPSC clones, and experiments. Here, we report the rapid (5 weeks) and efficient (~90%) induction of induced dopaminergic neurons (iDANs) through transient overexpression of lineage-promoting transcription factors combined with stringent selection across five donors. We observe maturation-dependent increase in dopamine synthesis, together with electrophysiological properties consistent with midbrain dopaminergic neuron identity, such as slow-rising after hyperpolarization potentials, an action potential duration of ~3ms, tonic sub-threshold oscillatory activity, and spontaneous burst firing at frequency of ~1.0-1.75 Hz. Transcriptome analysis reveals robust expression of genes involved in fetal midbrain dopaminergic neuron identity. Specifically expressed genes in iDANs, relative to their isogenic glutamatergic and GABAergic counterparts, were linked to the genetic risk architecture of a broad range of psychiatric traits, with iDANs showing particularly strong enrichment in loci conferring heritability for cannabis use disorder, schizophrenia, and bipolar disorder. Therefore, iDANs provide a critical tool for modeling midbrain dopaminergic neuron development and dysfunction in psychiatric disease.

## INTRODUCTION

Dopaminergic neurotransmission regulates human behavior, motivation, affect, and cognition^1^. Dysfunction of dopaminergic neurons is importantly involved in the pathogenesis of neurological and psychiatric disorders such as Parkinson disease^2^, substance use disorders^3^, and psychosis^4^.

Human induced pluripotent stem cell (hiPSC) models provide an approach to generate large numbers of disease-relevant cell types and investigate disease processes at the cellular and molecular level^5^, enabling the functional characterization of disease risk factors through genetic, pharmacologic, and physiological manipulations not possible in the relevant *in vivo* tissues^6,7^. Current methods to generate dopaminergic neurons *in vitro* either recapitulate key aspects of neurodevelopment through sequential application of small molecules and growth factors^8^ or overexpress exogenous transcription factors known to induce dopaminergic neuron identity^9–13^. A remaining limitation of current techniques is the high degree of variable reproducibility between hiPSC donor lines and investigator groups. The most widely utilized technique^8^ employs sequential additional of small molecules and protein factor combinations to the media based upon known developmental pathways; however, studies have reported highly inconsistent yields ranging from ~8% to >90%^14–18^. Consequently, many differentiations result in heterogenous cell populations^19,20^ and also include non-dopaminergic cells that are poorly characterized and of unknown relevance to the model system.

Here, we report the reliable induction of dopaminergic neurons from hiPSCs by transient overexpression of *ASCL1, LMX1B,* and *NURR1*^11^ (ALN) combined with antibiotic selection via a single doxycycline-inducible lentiviral vector, achieving a median percent purity across five independent donors of 92%. These induced dopaminergic neurons (<u>iDANs</u>) express genes consistent with midbrain regional patterning and dopaminergic neuron identity, demonstrate maturation-dependent dopamine synthesis, and electrophysiological hallmarks of *in vivo* dopaminergic neuron activity. Transcriptomic analyses of iDANs and post-mortem midbrain tissues provide further evidence of a fetal midbrain dopaminergic neuron identity of iDANs. Finally, specifically expressed genes in iDANs and isogenic induced GABAergic and glutamatergic neurons uncovered enrichment in risk loci for several psychiatric disorders, with evidence of neuronal subtype- and disorder-specific enrichments among biologically relevant pathways.

## MATERIALS AND METHODS

### Human induced pluripotent stem cell culture

All hiPSCs were derived by sendai viral OKSM reprogramming of dermal fibroblasts obtained from control donors in a previous cohort^21^. hiPSCs were maintained in StemFlex media (Gibco, #A3349401) supplemented with Antibiotic-Antimycotic (Gibco, #15240062) on Matrigel-coated (Corning, #354230) plates and passaged at 80−90% confluency with 0.5mM EDTA (Life Technologies, #15575-020) every 4-7 days for a maximum of 10 passages; no hiPSCs were cultured beyond passage 30. Routine cytogenetic analysis at WiCell confirmed normal karyotype of all donor lines. Donor meta-data are included in <u>Supplementary Table 1</u>.

### TetO-ALN-PuroR vector

We cloned a puromycin-resistance gene (*PuroR*) into a publicly available (Addgene: 43918) lentivirus vector encoding *TetO*-*ASCL1-LMX1B-NURR1-PuroR* (*“ALN”*). *PuroR* was inserted at the 3’ end of *NURR1* and separated by a 2A peptide sequence.

### Lentivirus production

Third-generation lentiviruses for *pUBIQ-rtTA* (Addgene 20342)*, tetO-ASCL1-LMX1B-NURR1-PuroR* (Addgene, 43918), *tetO-ASCL1-PuroR* (Addgene 97329)*, tetO-DLX2-HygroR* (Addgene 97330), and *tetO-Ngn2-PuroR-GFP* (Addgene 79823) were generated via polyethylenimine (PEI, Polysciences, #23966-2)-mediated transfection of human embryonic kidney 293T (HEK293T) cells using existing protocols^22^.

### Production of induced dopaminergic neurons (iDANs)

hiPSCs were harvested via incubation in Accutase Cell Detachment Solution (Innovative Cell Technologies, #AT104), quenched with DMEM (Gibco, #11965092), and centrifuged at RT at 800g for 5 minutes. Cell pellets were gently resuspended in StemFlex (Gibco, #A334901) supplemented with 10uM ROCK Inhibitor (StemCell Technologies, #72307) and counted with a Countess machine from Thermo Fisher Scientific (#AMQAX1000); the proportion of living cells was estimated by exclusion of Trypan Blue Solution, 0.4% (Gibco, #15250061). The cell suspension was then diluted in StemFlex with ROCK inhibitor to a concentration of 1e6 cell/mL and mixed with 50uL aliquots of both *tetO-ALN-PuroR* and *pUBIQ*-*rtTA* viruses tittered at an estimated 1 × 10^7^ IU/mL using a qPCR Lentivirus Titration Kit (Applied Biological Materials, #LV900). hiPSCs were plated on Matrigel-coated plates and incubated at 37°C with virus overnight. The following day, DIV1, the media was aspirated and replaced with Induction Media (see <u>Supplementary Note 1</u>). 1.0ug/mL puromycin (Sigma, #7255) was added the following day on DIV2. Media was changed on DIV3 if substantial cell death was present. Beginning on DIV5, media consisted of Induction Media with 1.0mg/mL puromycin with 2.0uM arabinosylcytosine (Sigma, #C6645) (“Ara-C”) to inhibit the proliferation of non-neuronal cells. On DIV7, cells were dissociated and replated in Induction Media supplemented with 1.0ug/mL doxycycline, 1.0ug/mL puromycin, 2.0uM Ara-C, and 10uM ROCK inhibitor on plates double-coated with 0.1% polyethylenimine (PEI) and 80ug/mL Matrigel. The following day (DIV8), media was replaced with Induction Media supplemented with 1.0ug/mL doxycycline and 2.0uM Ara-C. Ara-C was continued until DIV9, and doxycycline until DIV14, at which time the media was switched to Neuron Media. Half media changes were made every other day until the time of harvest (DIV35 for RNAseq). See <u>Supplementary Note 1</u> for media recipes, a more detailed protocol, and trouble-shooting information.

### Production of Induced GABAergic Neurons (iGANs)

iGANs were generated from two hiPSC donors (C-1 and C-2) via transduction with two separate doxycycline-inducible lentivirus vectors encoding *ASCL1-PuroR* and *DLX2-HygroR* according to Yang et al., 2017^23^, with slight modification; also see detailed method available in *Protocol Exchange*^24^. In brief, hiPSCs were harvested in Accutase, dissociated into a single-cell solution, quenched in DMEM, pelleted via centrifugation for 5 minutes at 800g, and resuspended in StemFlex with 10uM ROCK Inhibitor Y-27632. Volumetric equivalents of *ASCL1-PuroR, DLX2-HygroR,* and *pUBIQ-rtTA* were added to the suspension, mixed gently by inversion, dispensed onto Matrigel-coated plates, and incubated overnight at 37°C. The next day, media was changed to Induction Media (identical recipe to that used for iDAN generation) with 1.0μg/mL doxycycline (DIV1). 1.0μg/mL puromycin and 250μg/mL hygromycin were added the next day (DIV2) and continued for four days. We included 4.0μM Ara-C in the media from DIV4-8. Cells were harvested, dissociated, and replated on 0.1%PEI and 80ug/mL Matrigel-coated plates around DIV5-7. Media was switched to Neuron Media on DIV14, and doxycycline was withdrawn at that time. Half media changes were performed every other day from DIV14 until the time of harvest at DIV42 for the samples used for RNA-seq library generation.

### Production of Induced Glutamatergic Neurons (iGLUTs)

iGLUTs were generated via transduction of hiPSCs (donors C-1 and C-2) with *NGN2-eGFP-PuroR*^25,26^ using the same protocol steps used to produce iGANs, with the exception that cells were dissociated and replated on DIV3 or DIV4 and matured until DIV21 at which time they were harvested for RNA extraction and RNAseq library generation.

### RNA Extraction, Purification, and Quantification

Media was aspirated, and the cells were washed twice with PBS. Samples were lysed with TRIzol Reagent (Thermo, #15596026). RNA was extracted and purified using the Direct-zol RNA miniprep kit with in-column DNAse treatment (Zymo Research, #R2051). Purified RNA was eluted in UltraPure water and stored at −80°C until needed for rt-qPCR or RNAseq library preparation. RNA concentration was determined by running the samples on a Qubit 3 Fluorometer (Invitrogen, #Q33216) with the Qubit RNA HS Assay Kit (Thermo, #Q32852).

### Reverse-transcription quantitative qPCR (rt-qPCR)

50ng of RNA per each sample was loaded into a 384 plate and quantified using the *Power* SYBR Green RNA-to-C_t_ *1-Step* Kit (Thermo, #4389986). Reverse transcription and quantitative PCR took place on a QuantStudio 5 Real-Time PCR System (Thermo, #28570). Forward and reverse primer sequences for each gene are provided in <u>Supplementary Table 2</u>. Transcript abundance levels were quantified using the ΔΔ-C_t_ method^27^, with normalization of RNA input to *ACTB* as a loading control. For each gene and timepoint shown in **Figure 2,** data from samples generated from multiple donors were pooled to better capture any donor- and batch-related variance in the true expression value of the gene. For each gene, a one-way ANOVA with Tukey post-hoc testing was utilized to test for differences in expression level at each of the three time points (DIV0 (hiPSCs), DIV14, and DIV35) using the *aov* and *TukeyHSD* functions in R.

### Immunocytochemistry

We adapted a protocol from one of our previous reports^28^. At DIV7, immature iDANs were split onto glass coverslips in a 24-well plate and matured until the desired timepoint. At the time of harvest, the media was aspirated from each well followed by two washes with PBS. 500uL of 4% para-formaldehyde solution (Electron Microscopy Sciences, #15170) in PBS was added to each well and incubated at room temperature for 10 minutes followed by three PBS washes. 500uL of a blocking solution consisting of PBS with 5% donkey serum (Jackson, #017-000-121) and 0.1% Triton X-100 (Sigma, #T8787) was added to the wells and incubated for one hour at room temperature. After two PBS washes, diluted primary antibodies were added in 5% donkey serum and 0.1% Tween-20 (Boston BioProducts, #IBB-181X) in PBS and incubated overnight at 4 °C. The solution was then removed, and the wells were washed three times with PBS, followed by addition of secondary antibodies diluted in PBS and incubation in the dark for two hours. Finally, the secondary antibody solution was removed, the wells were washed three times with PBS, and 500μL of 0.5μg/mL DAPI (Sigma, #D9542) in PBS was added to the wells for a ten-minute incubation to stain cellular nuclei. Coverslips were carefully transferred to glass slides (Fisher Scientific, #12-544-7) and fixated using AquaPolymount (Polysciences Inc., #18606-20). See <u>Supplementary Table 3</u> for primary and second antibodies. The percent of cells positive for TH was determined by manual counting of all DAPI+ nuclei that also stained TH+ in random views of confocal images (containing at least 15 cells) taken across two or more separate experiments among five independent donors.

### Post-Mortem Brain Sample Preparation and Fluorescence-Activated Nuclear Sorting

Post-mortem sample processing, dissection, and fluorescence-activated nuclei sorting (FANS) haven been described previously^29^. In brief, substantia nigra pars compacta (SNpc) along with surrounding regions of the ventral tegmental area (VTA) were dissected from adult brains with a post-mortem interval of less than 24 hours. All donors were controls in an ongoing cohort study and did not have any known psychiatric illnesses. Frozen, but not fixed, tissue samples were homogenized in ice-cold lysis buffer and the resulting homogenate was mixed with a sucrose solution and ultra-centrifuged for one hour. The pellet was then resuspended and incubated with primary antibodies targeting NeuN (EMD Millipore, MAB377X; pre-conjugated with Alexa 488) and Nurr1 (Santa Cruz Biotechnology, sc-990), the latter of which had been incubated with Alexa Fluor 647 fluorochrome (Thermo Fisher, A27040) for one hour prior. After incubating in primary antibodies for two hours and DAPI (4′,6-diamidine-2′-phenylindole dihydrochloride, Sigma Aldrich, 10,236,276,001 Roche) for the last ten minutes, the nuclei suspension was processed on a FACSAria flow cytometry sorter. Donor meta-data are included in <u>Supplementary Table 1</u>.

### Nuclear RNAseq of Post-Mortem Midbrain Samples

Nuclear RNAseq libraries from NeuN+/Nurr1+ and NeuN-/Nurr1-nuclei were prepared as described previously^29^. In summary, nuclei were lysed in TRIzol LS Reagent (Thermo Fisher, #10296028) and mixed with an equal volume of 100% ethanol. DNA-digestion and RNA-extraction was performed using the Zymo-Spin IC Column from Direct-zol RNA MicroPrep kit (Zymo Research, R2060) per the manufacturer’s instructions. The yield and quality of the resulting RNA samples were assessed with the Agilent Bioanalyzer using an Agilent RNA 6000 Pico kit (Agilent, #5067-1513). Ribosomal rRNA-depleted RNAseq libraries were prepared using the SMARTer Stranded RNA-Seq kit (Clontech, #634836) according to the manufacturer’s instructions with the following specifications: (a) RNA was fragmented at 94 °C for three minutes; (b) after index annealing with the Illumina indexing primer set (Illumina, #20020492), 12 PCR cycles were used for cDNA amplification. Libraries were subsequently purified using a 1:1 volumetric ratio of SPRI beads (Beckman Coulter Life Sciences, #B23318) to remove primer dimers and enrich for a target library size of ~300bp, which was confirmed on the Agilent Bioanalyzer.

### Whole-Cell RNAseq Library Preparation and Sequencing of In Vitro Samples

Strand-specific, rRNA-depleted RNA-seq libraries were prepared from 100-1,000ng RNA per sample using the KAPA RNA HyperPrep Kit with RiboErase (HMR) (Roche, #KK8560). RNA fragmentation was performed at 94°C for 6 minutes and 10 PCR cycles were used during library amplification with TruSeq single-index adapters (Illumina, #20020492). Final library concentrations were quantified with both Qubit fluorometric quantification (DNA dsDNA HS kit, Thermo, #Q32851) and the KAPA Library Quantification kit (Kapa Biosystems, #KK4873). The samples were run on an Agilent 2100 Bioanalyzer (Agilent #G2939BA) with the High Sensitivity DNA Kit (Agilent, #5067-4626) to confirm the appropriate distribution of fragment sizes and the absence of significant artifactual contaminants. 150 base-pair paired-end sequencing was performed on a NovaSeq 6000 System to a desired depth of ~50,000,000 reads per library.

### RNA-sequencing analysis

a. *<u>Library-processing</u>*: raw data were aligned to the genome (GRCh38) using *STAR* v2.7.0^30^. Counts per gene were extracted using the *featureCounts*^31^ function in the *Rsubread* package^32^. Sample count matrices were processed in R version 4.0.2 using the *limma*^33^ and *edgeR*^34,35^ packages. For the dataset consisting of iDANs, hiPSCs, midbrain NeuN+/Nurr1+ nuclei (midbrain dopaminergic neurons, or “MDNs”), and midbrain NeuN-/Nurr1-nuclei (“non-MDNs”), lowly expressed genes were filtered out using the *filterByExpr* function in *edgeR*, leading to a reduction in the total number of ensemble gene IDs from 58, 037 to 16,641. Library log_2_(CPM) distributions were normalized with the trimmed mean of M-values method^36^. Heteroscedasticity was removed from the data using the *voom* function of *limma* and linear models were fit for cell type-contrasts of interest with weights generated using empirical Bayes moderation^37^. The results of library-processing and quality controls are shown in <u>Supplementary Figure 1</u>.
b. *<u>Derivation of differentially and specifically expression genes</u>*:“Differentially expressed genes (DEGs)” were defined as genes with a log_2_(fold change in CPM) of at least 1.0 and that surpassed a significance threshold of p < 0.05 after correcting for multiple testing with the Benjamini-Hochberg procedure in the *decideTests* function in *limma*. The in vitro RNA-seq dataset consisting of hiPSCs, iDANs, iGANs, and iGLUTs was processed similarly with the following two changes: (1) we did not filter out lowly expressed genes in order to capture all genes with potential specificity for a given cell type, and (2) we derived “specifically expressed genes (SEGs)”, instead of DEGs, by first contrasting the expression of all genes in each of the four cell types to the expression levels in the other three; SEGs were defined as genes in the top 10^th^ percent of *t*-statistics for each cell type (<u>Supplementary Table 4</u>), as in previous reports^38,39^. Genes with the highest 10% of t-statistics were deemed “specifically expressed,” with the caveat that there is indeed some degree of overlap in SEGs; accordingly, specifically expressed genes are not *exclusively* expressed in one cell type alone.
c. *<u>Gene-set overrepresentation analyses (GSOA)</u>*: We used *clusterProfiler* to perform GSOA on (i) DEGs in the dataset consisting of hiPSCs, iDANs, MDNs, and non-MDNs, and (ii) SEGs with enrichment in psychiatric disorder heritability. Tested gene sets included KEGG pathways^40^ and Gene Ontology Biological Processes^41,42^ (BP). We used an FDR threshold of < 0.05, and significance values are indicated throughout the text by the corresponding FDR q values. Significant results were visualized with tile plots, with the magnitude of the -log(FDR q) represented by the height of the tile. Enrichment maps showing relationships among pathways/processes based upon overlapping genes were created using the *emapplot* function in *clusterProfiler* after reducing the degree of redundancy in BP terms using semantic similarity analysis^43^ with *GOSemSim*^44^. Gene-concept network plots for the top enriched pathways were created with the *cnetplot* function of *clusterProfiler* to create gene networks.
d. *<u>Competitive gene-set testing for enrichment in cell type identity SEGs</u>*: a competitive gene-set testing procedure was conducted using Correlation-Adjusted Mean Rank gene-set test (CAMERA)^45^ on hiPSCs, iDANs, and the post-mortem tissue type DEGs to test for enrichment among specifically expressed genes in previously published datasets for brain cell types. We defined these brain cell type SEGs as those genes with the top 1% of specificity in the K1 and K2 cell types reported by Skene et al. (2018)^46^ and all of those reported by La Manno et al (2016)^20^ on developing midbrain cell types (<u>Supplementary Table 5</u>). The full results are shown without a specific FDR threshold cut off.
e. *<u>Correlation analyses of SEG enrichments in psychiatric risk loci</u>*: we calculated the spearman correlation coefficients in gene-level z-scores for those neuronal subtype-specific SEG sets with enrichment in any psychiatric disorder. A correlation matrix was generated in R using the *corr.test* function of the *psych* package, with adjustment of p values with Bonferroni correction; results with an adjusted p value < 0.05 were considered significant.

### Assessment of In Vitro Neuron Subtype Heritability Enrichment of Psychiatric Risk Loci

We intersected cell-type-specific expression patterns with genetic risk of 11 specified neurodevelopmental and neuropsychiatric disorders (attention-deficit/hyperactivity disorder^47^ (ADHD), anorexia nervosa^48^ (AN), autism spectrum disorder^49^ (ASD), alcohol use disorder^50^ (AUD), bipolar disorder^51^ (BIP), cannabis use disorder^52^ (CUD), major depressive disorder^53^ (MDD), obsessive-compulsive disorder^54^ (OCD), post-traumatic stress disorder^55^ (PTSD), and schizophrenia^56^ (SCZ), as well as Cross Disorder^57^ (CxD) GWAS summary statistics), along with Alzheimer’s disease^58^ (AD) and Parkinson’s disease^59^ (PD) to identify disorder-relevant cell types (<u>Supplementary Table 6</u>). We performed these cell-type association analyses using multi-marker analysis of genomic annotation (MAGMA)^60^. Four gene sets were defined by the protein-coding genes present in curated lists of SEGs for hiPSCs, iDANs, iGANs, and iGLUTs. Using MAGMA, SNPs were mapped to genes based on the corresponding build files for each GWA summary dataset. We ran gene analysis on GWAS summary statistics using the default method: snp-wise=mean (a test of the mean SNP association). A competitive gene set analysis was then used to test enrichment in genetic risk for a disorder across the four cell-type specific gene sets, with an adjusted p-value threshold of < 0.05.

### Multi-electrode array (MEA)

Commercially available human astrocytes (HA; Sciencell, #1800; isolated from fetal female brain) were thawed and seeded onto matrigel-coated 100 mm culture dish in commercial astrocyte medium (Sciencell, #1801) and expanded three passages in Astrocyte medium. Upon confluency, cells were detached, spun down and resuspended with Astrocyte medium supplemented with Antibiotic-Antimycotic (Anti/Anti; Thermo Fisher Scientific, #15240062) and split as 1 × 10^5^ cells per well on matrigel-coated 48 well CytoView MEA plates (Axion Biosystems). HAs were fed by full medium change with the Brainphys medium (2% FBS + Anti/Anti) + 2 μM Ara-C. At day 7, iDANs were split on the HAs with neuron media supplemented with 2% FBS by gently detaching them with Accutase for one hour, centrifuging (1000g × 5 mins), and resuspending in neuronal medium supplemented with 2% FBS and 5uM ROCK Inhibitor. After counting cells with a Countess machine, iDANs were seeded on the astrocyte culture (1 × 10^5^ cells/well). The media was changed the next day to neuronal medium supplemented with 0.5% FBS and 2 μM Ara-C. Half media changes were performed twice a week, one day before MEA measurement. 2 μM Ara-C treatment was discontinued after one week. MEA plates were measured twice a week on a Maestro Multi-electron array system (Axion Biosystems) at 37°C starting on day 21 of iDAN differentiation. For each measurement, plates were equilibrated in the machine for 5 min followed by 10 min recording, with spontaneous neural real-time configuration at threshold of 5.5. The plates were measured until week six of neuronal maturation followed by batch processing of files and analysis of compiled statistics.

### Electrophysiology

Neurons from two donors (C-1 and C-2) were plated on acid etched coverslips and co-cultured with human fetal astrocytes in Brainphys media to promote maturation^61^. Recordings were performed at five to six weeks after induction. Coverslips were transferred to a bath filled with modified aCSF solution, adapted from a mouse slice electrophysiology protocol^62^ containing NaCl 119 mM, D-glucose 11 mM, NaHCO_3_ 26.2 mM, KCl 2.5 mM, MgCl_2_ 1.3 mM, NaH_2_PO_4_ 1 mM, CaCl_2_ 2.5 mM (pH adjusted to 7.3 with HCl). Glass microelectrodes of 4.0 – 4.6 MΩ resistance were filled with an internal solution of 140 mM Potassium D-Gluconate, 4 mM NaCl, 2 mM MgCl_2_ × 6-H_2_O, 1.1 mM EGTA, 5 mM HEPES, 2 mM Na_2_ATP, 5mM NaCreatinePO_4_, and 0.6 mM Na_3_GTP. Chemicals obtained from Sigma-Aldrich. All solutions were ~ 295 mOsm. Whole-cell currents were recorded with an Axopatch 200B amplifier with application of manual series-resistance and capacitance compensation, filtered at 10 kHz for current-clamp and 1kHz for voltage-clamp, and digitized at 20 kHz and 10kHz, respectively, with the 1550 Digidata digitizer (Molecular Devices). For current-clamp recordings, a holding current was applied to set the resting potential to −65mV and 1 s current steps were applied in 0.02 nA increments. Spontaneous activity was measured in I=0 current-clamp mode. For voltage-clamp recordings, voltage steps (200 ms) were applied in 10 mV increments from a holding voltage of −80 mV to 50 mV. All voltage measurements were corrected for a calculated junction potential of −16.1 mV. Data were collected and analyzed using Molecular Devices pClamp 11 software and with custom-made routines written in R.

### Dopamine ELISA

For whole-cell dopamine ELISA, cells were harvested with Accutase and spun down for 5 minutes at room temperate at 1000g. The media supernatants were completely aspirated, and the cell pellets were flash frozen in liquid nitrogen. The ELISA was carried out using the Dopamine Research ELISA Kit from ALPCO (#17-DOPHU-E01-RES) according to the manufacturer’s instructions, beginning with cell lysis in homogenization solution in a Dounce homogenizer. Each sample was split into three technical triplicates. Absorbance at 450 nm was measured on a Varioskan LUX multielectrode microplate reader. A non-parametric local regression curve was fit to the values of the standards using the *loess* function in R, with the absorbance as the predictor variable and log_2_-transformed concentration (nM) as the response variable. Concentrations of the samples were extrapolated from the sample absorbances using the *predict* function in R using the regression model fitted to the standards.

## RESULTS

### Transient overexpression of ASCL1, LMX1B, and NURR1 induces dopaminergic neurons

Previous reports demonstrated that overexpression of *ASCL1*, *LMX1A*, and *NURR1* (also known as *NR4A2*) in human fibroblasts^11^ and hiPSCs^10^ could result in dopaminergic neurons with low yields limited to ~5% and ~33% purity, respectively. We designed an improved vector for induction of dopaminergic neurons (iDANs) from hiPSCs (**Fig. 1A**) that incorporated antibiotic selection (*TetO*-*ASCL1-LMX1B-NURR1-PuroR*, *“ALN-PuroR”*) (**Fig. 1B**). Doxycycline was administered until 14 days *in vitro* (DIV) (**Fig. 1C**), while selection with puromycin occurred from DIV2 to DIV6, and Ara-C was added from ~DIV4-6 to eliminate residual mitotic cells (see <u>Supplementary Note 1</u>). Early neuronal processes appeared at DIV7, and iDANs were matured for 35 days, at which time they show extensive branching and lengthy processes (**Fig. 1D**).

**Figure 1:**
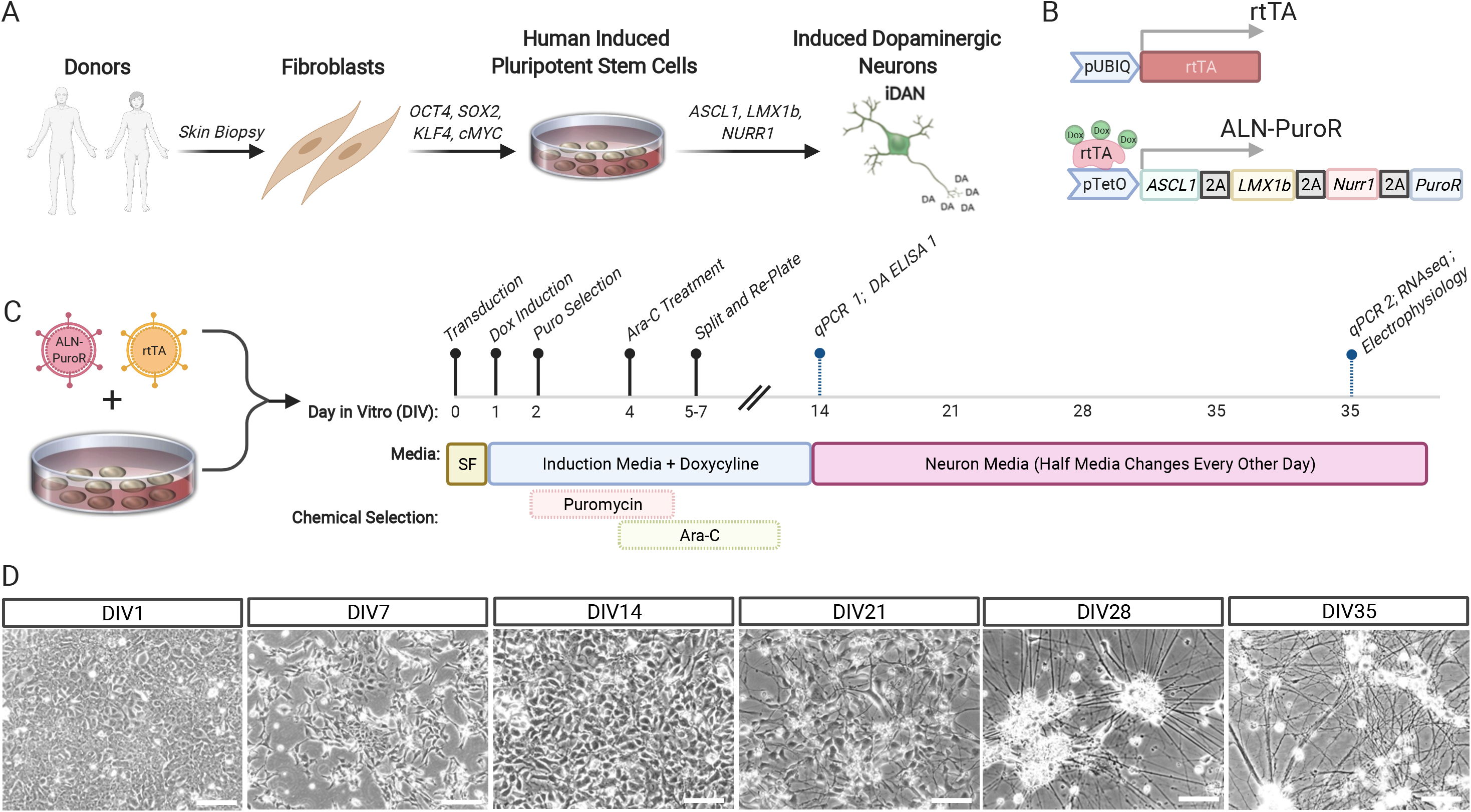
Production of induced dopaminergic neurons with *ASCL1, LMX1B,* and *NURR1* transduction. (A) Schematic showing overall process of producing iDANs from hiPSCs. (B) Cartoon illustrating key features of the *tetO*-*ALN-PuroR* and *rtTA* vectors. (C) Timeline of iDAN generation, beginning with transduction of hiPSCs at DIV0 and ending with sample harvesting; “SF” stands for StemFlex media. (D) Weekly brightfield images showing progressive development of neuronal morphology in iDANs. White scale bars = 50 μm.

Across five independent donor lines, qPCR revealed that DIV14 iDANs showed increased expression of the neuronal genes *MAP2* and *SYN1*, as well as *TH*, the rate-limiting enzyme in dopamine biosynthesis, and *AADC*, which converts L-DOPA to dopamine (**Fig. 2A**). They likewise showed robust expression of the midbrain marker genes *LMX1A*, *MSX1*, *EN2*, *FOXA2*, and *PITX3*, and a modest increase in expression of the dopamine transporter (*DAT*) and *VMAT2,* which did not reach statistical significance (**Fig. 2A**). By DIV35, *TH*, *LMX1A, OTX2, EN2,* and *FOXA2* had further increased (**Fig. 2A**). At the protein level, DIV35 iDANs co-expressed TH with MAP2 (**Fig. 2B**), SYN1 (**Fig. 2C**), NEUN (**Fig. 2F**), DAT (**Fig. 2E**), as well as the midbrain marker OTX2 (**Fig. 2D**) and GIRK2, an inwardly rectifying potassium channel that mediates D2R stimulation^63^ (**Fig. 2E**).

**Figure 2:**
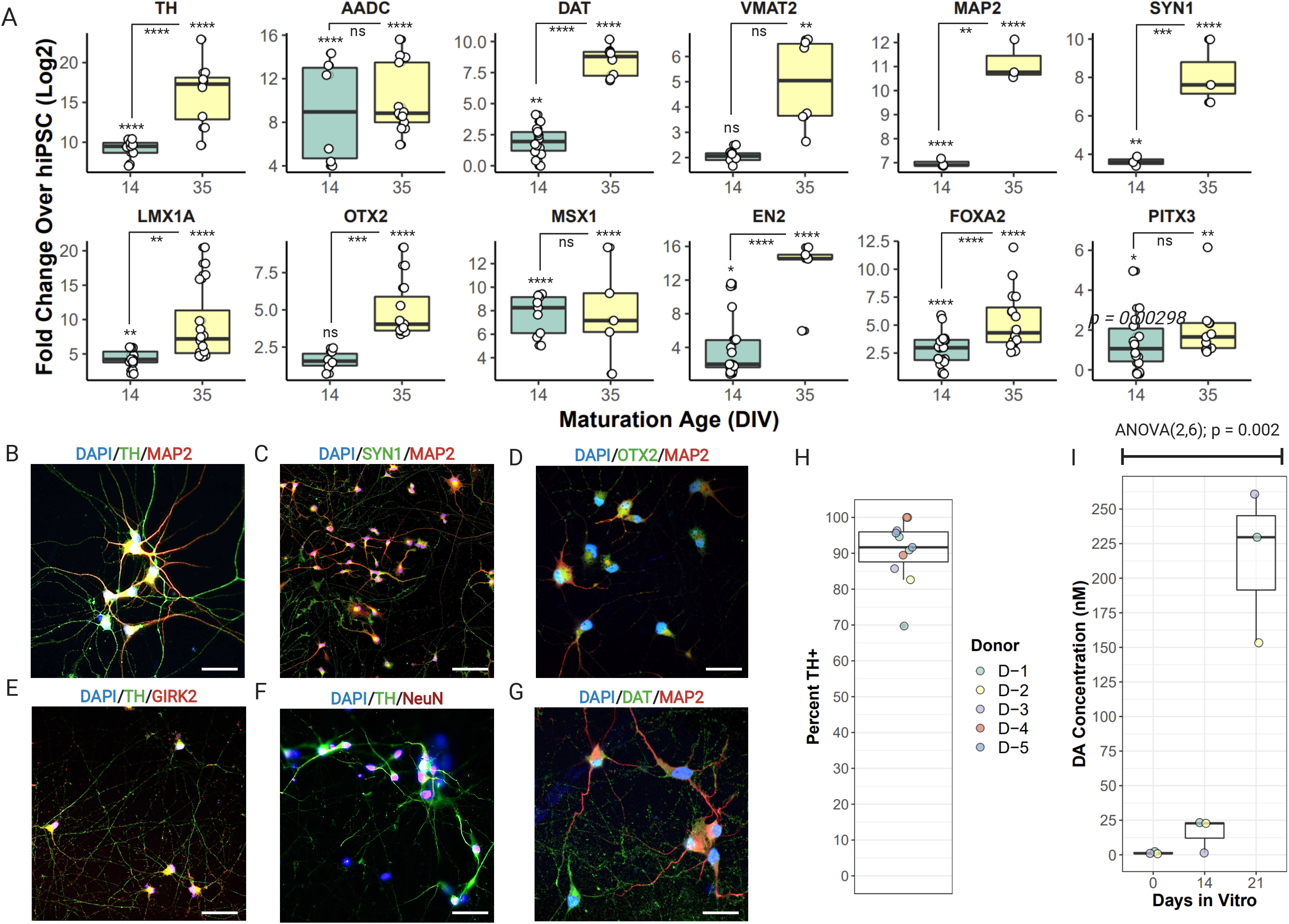
Marker gene expression, purity, and dopamine production in iDANs. (A) Fold-change above hiPSCs in the expression of *TH, AADC, DAT, VMAT2, MAP2, SYN1, LMX1A, OTX2, MSX1, EN2, FOXA2,* and *PITX3*. A maturation-dependent increase in expression was seen for the majority of genes. Confocal images of immunocytochemical staining of DIV35 iDANs with (B) TH and MAP2, (C) SYN1 and MAP2, (D) OTX2 and MAP2, (E) TH and GIRK2, (F) TH and NEUN, and (G) DAT and MAP2. (H) Across five donors with replicate experiments, a median of 92% of all cells are TH+. (I) Maturation-dependent increase in dopamine biosynthesis, with all three independent donors tested showing dopamine production by DIV21. * p < 0.05, ** p < 0.01, *** p < 0.001, **** p < 0.0001, ns = not significant. DIV = “days *in vitro*”.

A median of ~92% of cells were positively stained for TH across five donors, with two or more replicate experiments per donor (**Fig. 2H)**. Moreover, ELISA analysis of iDANs from three donors confirmed a maturation-dependent increase in total dopamine biosynthesis (ANOVA (2,6) = 20.78; p = 0.0020); by DIV21, dopamine production across the three donors was significantly elevated relative to DIV0 (p = 0.0017) and DIV14 (p = 0.018) (**Fig. 2I**). Altogether, transduction with *ALN-PuroR* leads to robust induction of >90% dopaminergic neurons (range: 70-98%), showing widespread dopaminergic marker gene expression and dopamine biosynthesis in a maturation-dependent manner.

### iDANs show physiological hallmarks of in vivo dopaminergic neurons

Across two independent donors, multi-electrode array recordings revealed increasing burst frequency (Hz), weighted mean firing rate (Hz), network burst frequency (Hz), the fraction of active electrodes with bursting activity, and coefficient of variation of the inter-spike interval (ISI, a measure of maturation age^64^) across maturation (<u>Supplementary Figure 2</u>). In contrast, burst duration, network spike duration, and spikes per network burst remained steady across maturation, in agreement with previous MEA analyses of developing primary rodent cortical neurons in vitro^64^.

With the same two donors, we examined the intrinsic excitability of iDANs using patch-clamp electrophysiology. After approximately five weeks of induction, iDANs exhibited regenerative action potentials in response to current injections (**Fig. 3A**), with a notable slow after-hyperpolarization potential (AHP) (**Fig. 3B**) typical of dopaminergic neurons^65^. The action potential width of 3 ms was similar to that reported for primate/rodent DA neurons^66^. We also observed prominent voltage-gated sodium and potassium currents but not an I_h_ inward current (**Fig. 3C**). The cell capacitance for iDANs (20 ± 8 pF) was smaller than in rodents^67^. iDANs exhibited spontaneous activity at resting membrane potentials (**Fig. 3E**), with some showing continuous tonic-like firing (**Fig. 3D**). We compared the distributions of spontaneous firing for both patch-clamp and MEA assays and observed a median frequency of about 1.0 – 1.75 Hz (**Fig. 3F, G**). The basic neuronal properties (e.g., capacitance, resting potential) and firing behavior were indistinguishable between the two donor lines, highlighting the replicability of this induction method across individuals (<u>Supplementary Table 7</u>). Altogether, iDANs displayed many of the electrophysiological hallmarks of their *in vivo* midbrain dopaminergic neuron counterparts.

**Figure 3:**
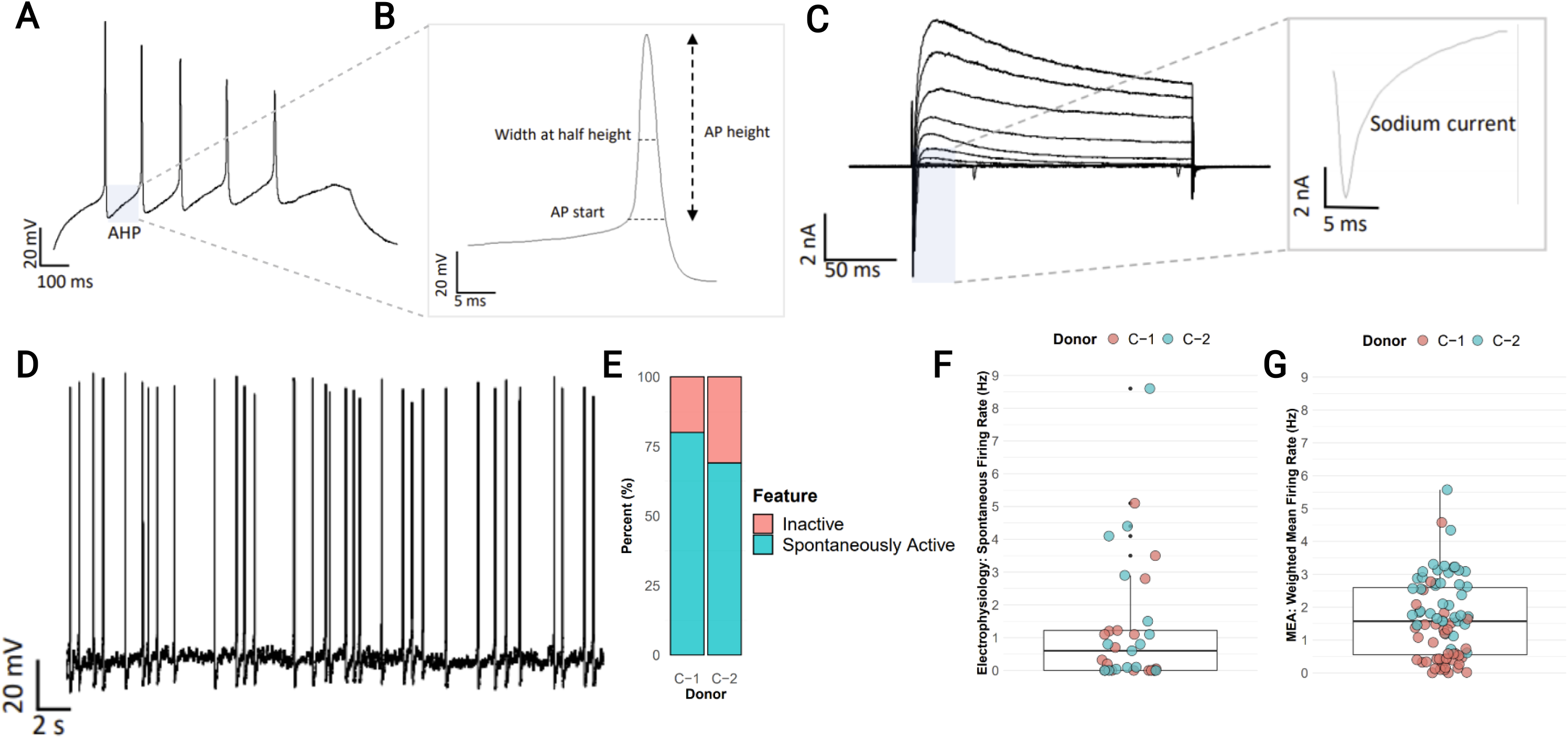
Electrophysiological characterization of iDANs. (A) A representative voltage trace shows evoked action potentials with current injection step (.02 pA). Highlighted area (grey box) illustrates the slow after-hyperpolarization potential (AHP). (B) Enlarged view of a representative iDAN action potential, illustrating threshold, height, and duration (width at half-height) measurements. (C) Representative traces of voltage-gated potassium and sodium currents evoked by voltage steps from −80 mV to 50 mV. Inset shows inward sodium current at −50 mV. (D) Example of spontaneous tonic firing at resting potential (V_m_ = −53 mV). (E) Proportion of spontaneously active neurons by donor (N=18 C-1, N=15 C-2). (F,G) Comparison of spontaneous firing rates measured by whole-cell patch-clamp with a mean frequency of 1.28 Hz (F, cells (n)= 18,15 for C-1 and C-2, respectively) and by multielectrode array (MEA), with a median weighted mean firing rate (WMFR) of 1.64 (G, wells (n) = 39, 50).

### iDANs exhibit a fetal midbrain dopaminergic neuron transcriptomic profile

To benchmark iDANs to a reference *in vivo* dataset, we conducted an RNAseq analysis of neurons and non-neuronal cells sorted from post-mortem midbrain (**Fig. 4A**), comparing midbrain dopaminergic neurons (NeuN+/Nurr1+, nuclei, “MDNs”) and midbrain non-dopaminergic neurons (NeuN-/Nurr1-, “Non-MDNs”). Principal component analysis of total gene expression across all samples revealed distinct clustering by cell type, with iDANs aligning with MDNs on PC1, which accounted for 72% of the total variance (**Fig. 4B**). Hierarchical clustering separated hiPSCs and iDANs from the post-mortem samples but also demonstrated greater relatedness to MDNs than non-MDNs (**Fig. 4C**).

**Figure 4:**
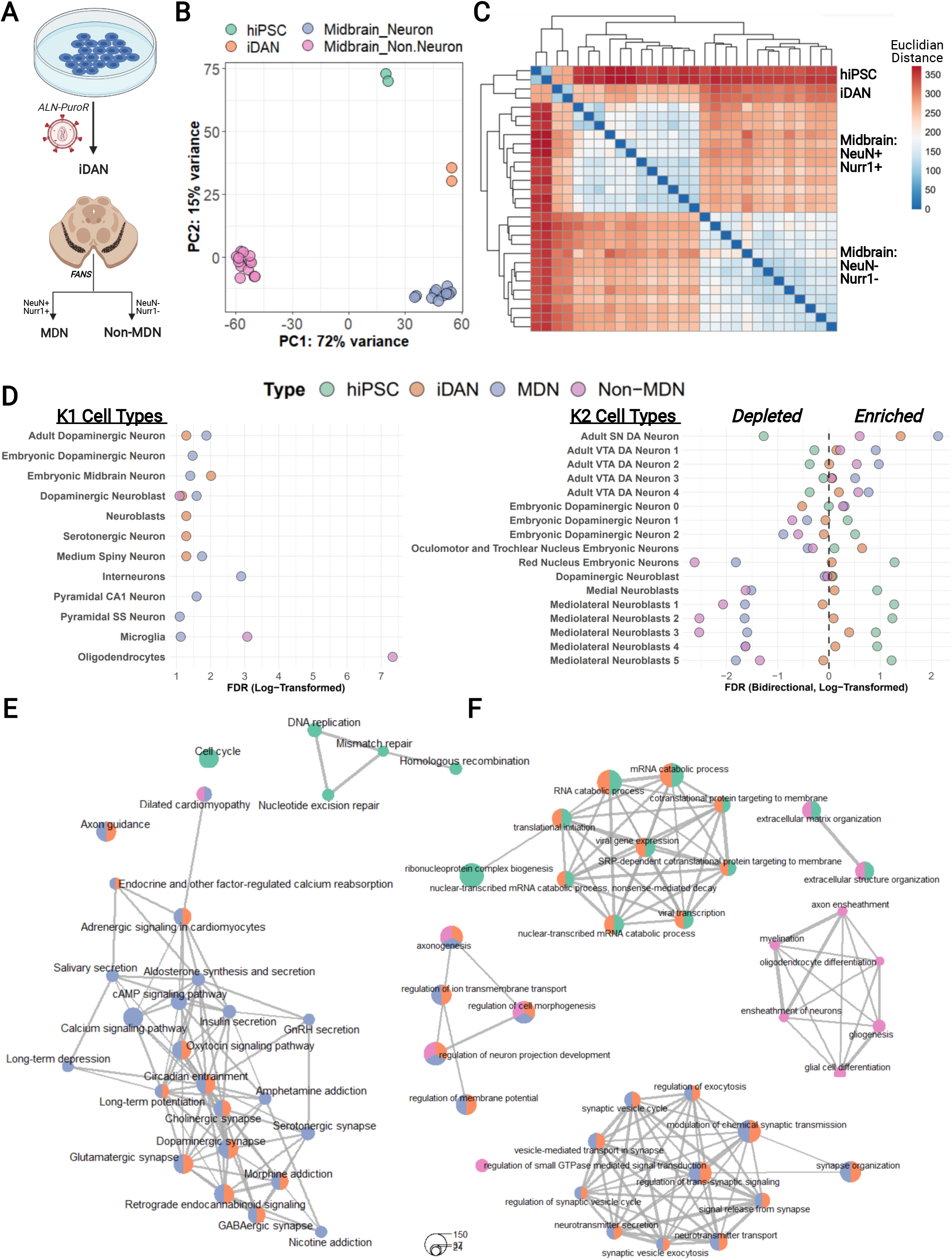
Transcriptomic analysis of iDANs. (A) Schematic showing the generation iDANs and post-mortem samples with fluorescence-activated nuclear sorting from human midbrain. (B) Principal component analysis on PC1 and PC2 shows strong clustering of samples by cell type, with iDANs aligning with post-mortem midbrain dopamine neurons (MDNs) on PC1. (C) Hierarchical clustering of RNA-seq samples by Euclidean distances between transcriptomic profiles (all 16, 641 genes), revealing sample relatedness by cell type. (D) Results of competitive gene set testing for enrichment in specifically expressed genes from the K1 (left) and K2 (right) cell types from Skene et al., 2018^46^. Left: for the K1 results, hiPSC data points are omitted for graphical purposes, and only enrichments in the positive direction are shown; full results are contained in <u>Supplementary Figure 3</u>. Right: selected results for dopaminergic lineage cell types from the K2 datasets, with enrichments in the “down” direction represented by log-transformed FDR q values multiplied by −1. The full results including all 149 K2 cell types are shown in <u>Supplementary Figure 4</u>. (E) Enrichment maps depicting top KEGG pathways enriched among cell-type-specific gene expression. Individual pathways are shown as circular “nodes”, with the node color indicating cell type and node size representing the number of genes within the pathway node overlapping the specifically expressed genes for that cell type. Pathway nodes are connected by edges to form networks based upon overlapping genes. Spaces between pathway networks and free-standing nodes are not biologically meaningful, as individual networks were arranged for graphical purposes. (F) Same as in E, but with nodes representing Gene Ontology Biological Processes instead of KEGG pathways.

We generated DEGs from our *in vitro* and *in vivo* cell types and performed a competitive gene set testing procedure to explore potential enrichments in established brain cell-type-specific marker genes (**Fig. 4D**, <u>Supplementary Figure 3</u>). Among a group of 24 cell-type-specific marker gene lists^46^ (<u>Supplementary Table 5</u>), MDNs showed strong enrichment in the positive direction for several cell types including “Interneuron” (FDR q = 1.27 × 10^−3^) and “Adult Dopaminergic Neuron” (FDR q = 0.0133). While the magnitude of iDAN enrichment in “Adult Dopaminergic Neuron” (FDR q = 0.0516) was somewhat lower than that observed for MDNs, iDANs were most highly enriched in “Embryonic Midbrain Neurons” (FDR q = 9.69 × 10^−3^). In contrast, the non-MDNs showed high enrichment in the gene sets specific to “Oligodendrocytes” (FDR q = 4.68 × 10^−8^) and “Microglia” (FDR q = 8.42 × 10^−4^). We then expanded the analysis to a more refined set of 149 specific cell types^46^ (<u>Supplementary Table 5</u>), each of which belong to one of the 24 broader cell type classifications in the first dataset (<u>Supplementary Figure 4</u>). Among the cell types belonging to the dopaminergic neuron lineage, both MDNs (FDR q = 7.38 × 10^−3^) and iDANs (FDR q = 0.0398) were most enriched in “Adult Substantia Nigra Neurons”, with iDANs showing additional positive enrichment in early developmental cell types (<u>Supplementary Table 9</u>). With a third dataset^20^, this time derived from early developmental midbrain cell types (<u>Supplementary Table 5</u>), competitive gene set testing confirmed that iDANs were most strongly enriched in specifically expressed genes defining early midbrain and dopaminergic neurons and progenitor cells (<u>Supplementary Figure 5</u>).

**Figure 5:**
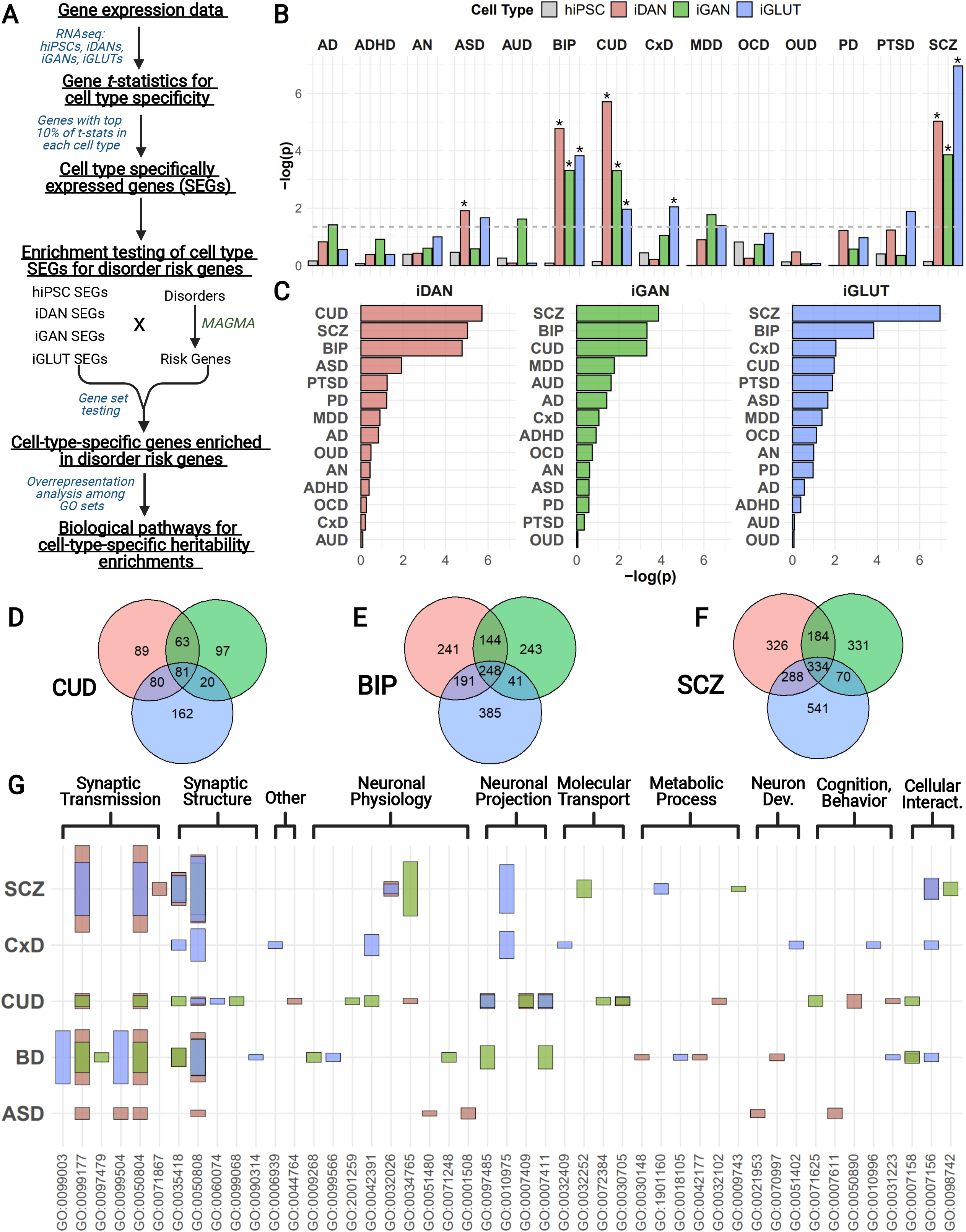
Neuronal subtype heritability enrichment for psychiatric disorders among hiPSC-derived neurons. (A) Analysis workflow for identification of biological pathways implicated in neuronal subtype-specific heritability enrichments in psychiatric disorders^38^. (B) Gene set testing results utilizing MAGMA-derived risk genes and in vitro cell type SEGs for hiPSCs, iDANs, iGANs, and iGLUTs. Dashed line indicates nominal significance (p < 0.05), while * indicates those SEG sets that were significantly enriched after correction for multiple testing. Alzheimer disease = AD; attention-deficit/hyperactivity disorder = ADHD; anorexia nervosa = AN; autism spectrum disorder = ASD; alcohol use disorder = AUD; bipolar disorder = BIP; cannabis use disorder = CUD; cross-disorder pleiotropic loci = CxD; major depressive disorder = MDD; obsessive-compulsive disorder = OCD; opioid use disorder = OUD; Parkinson disease = PD; post-traumatic stress disorder = PTSD; schizophrenia = SCZ. (C) Plot showing the results of MAGMA by each cell type group. Overlap among significantly enriched neuronal subtype-specific genes among (D) CUD, (E) BIP, and (F) SCZ. (G) Results from gene set overrepresentation analyses of significantly enriched gene sets and their involvement in biological processes. GO sets are grouped into meta categories; the set names corresponding to each GO ID number are listed in the full results in <u>Supplementary Table 11</u>. Bar colors indicate neuronal subtype, and bar height represents the magnitude-log(FDR q values).

Finally, we conducted gene set overrepresentation analyses (GSOA) to evaluate the biological relevance of cell-type-specific gene expression. Broadly, enriched terms were consistent with the known identities and functions of the respective cell type (<u>Supplementary Table 8</u>). While hiPSC gene expression was enriched in KEGG pathways^40^ involved in the cell cycle (e.g., “Cell Cycle”, q = 4.08 × 10^−10^), both MDNs and iDANs were enriched in pathways with clear links to dopaminergic neuron biology, such as “Dopaminergic Synapse” (q = 0.000163 in iDANs; q = 7.08 × 10^−11^ in MDNs), “Long-Term Potentiation” (q = 0.030 in iDANs; q = 2.16 × 10^−6^ in MDNs) and “Morphine Addiction” (q = 0.010 in iDANs; q = 3.41 × 10^−10^ in MDNs) (**Fig. 4E**). These findings were corroborated by additional enrichment in a network of Gene Ontology Biological Processes^41,42^ related to synaptic structure (e.g., “Synapse Organization” (q = 1.30 × 10^−21^ in iDANs; q = 3.54 × 10^−21^ in MDNs)) and neurotransmission (e.g., “Dopamine Secretion” (q = 3.16 × 10^−6^ in iDANs; q = 0.0060 in MDNs)), whereas non-MDNs were enriched in processes pertaining to glial cells (e.g., “Glial Cell Differentiation”, q = 1.49 × 10^−8^; “Myelination”, q = 1.71 × 10^−6^) (**Fig 4F**). in total, these results support a fetal-like midbrain dopaminergic neuron identity for iDANs.

### Differential enrichment of induced dopaminergic, GABAergic, and glutamatergic neurons in psychiatric risk genes

We next sought to test whether hiPSC-derived neuronal subtype-specific gene expression was enriched in psychiatric disease risk loci. After generating isogenic iDANs, iGANs, and iGLUTs and calculating those SEGs most specifically expressed in each induced neuronal subtype, we applied (MAGMA)^60^ to test for the enrichment of *in vitro* cell type SEGs among an array of psychiatric disorders^47–56,68^ (**Fig. 5A**), as well as a set of pleiotropic loci implicated in a cross-disorder (CxD) analysis of eight psychiatric conditions^57^. Alzheimer’s disease^58^ (AD) and Parkinson’s disease^59^ (PD) were included as brain-related but non-psychiatric disorders. SEGs for iDANs, iGANs, and iGLUTs were significantly enriched in risk genes for cannabis use disorder (iDAN: p = 1.94 × 10^−6^; iGAN: p = 4.89 × 10^−4^; iGLUT: p = 0.0109), bipolar disorder (iDAN: p = 1.68 × 10^−5^, iGAN: p = 4.82 × 10^−4^; iGLUT: p = 1.48 × 10^−4^), and schizophrenia (iDAN: p = 9.32 × 10^−6^, iGAN: p = 1.37 × 10^−4^; iGLUT: p = 1.10 × 10^−7^), with iDANs showing additional enrichment in autism spectrum disorder (p = 0.0122) and iGLUTs showing enrichment in the cross-disorder pleiotropic loci (p = 0.00896) **(Fig 5B**). As expected, hiPSCs were not enriched in any condition (<u>Supplementary Table 10</u>). Overall, iDANs were most enriched in CUD, while both iGANs and iGLUTs were most enriched in SCZ (**Fig. 5C**), consistent with emerging data that glutamatergic and GABAergic neurons are particularly impacted by SCZ genetic risk loci^46,69–71^. Further supporting our findings of heritability enrichment for iDAN SEGs, we also found that SEGs from our post-mortem MDNs were similarly enriched in risk genes for BIP (p = 5.82 × 10^−6^), SCZ (p = 0.00074), and CUD (0.016), along with an additional enrichment in ADHD (p = 0.012) (<u>Supplementary Figure 6</u>).

For each disorder with enrichment among one or more hiPSC-derived neuronal subtypes, we queried the biological relevance of the significantly enriched genes (<u>Supplementary Table 10</u>). Across all disorders and neuronal subtypes, we found consistent enrichment among biological processes related to synaptic transmission and structure (**Fig. 5G** and <u>Supplementary Table 11</u>); however, there was also significant neuronal subtype specificity among enriched pathways within the individual disorders. In SCZ, for example, only iDAN SEGs were overrepresented in processes such as “Monoamine Response” (q = 3.83 × 10^−3^) and “Response to Auditory Stimulus” (q = 1.41 × 10^−3^), while only iGAN SEGs were overrepresented in the term “Auditory Behavior” (q = 4.27 × 10^−3^), and iGLUT SEGs were uniquely overrepresented in the term “Cognition” (q = 5.13 × 10^−9^) (<u>Supplementary Figure 7</u>). Further illustration of differential enrichment of induced neuronal subtypes in BIP and CUD are shown in Supplementary Figures 8 and 9, respectively.

Having shown that unique subtypes of induced neurons are enriched in both shared and subtype-specific pathways *within* three major psychiatric disorders, we next assessed the extent to which gene expression specific to a given neuronal subtype is differentially enriched *across* disorders. Specifically expressed genes in iDANs demonstrated heritability enrichment in SCZ, BIP, CUD, and ASD (**Fig. 6A**). Intriguingly, iDAN SEGs enriched in ASD heritability, for example, were uniquely overrepresented in biological process such as “CNS Differentiation” (q = 7.31 × 10^−3^) and “Learning or Memory” (q = 1.31 × 10^−3^) (**Fig. 6B**), while those implicated in CUD risk alone formed a distinct pathway network related to neuron projection guidance (e.g., “Axon Guidance”, q = 8.08 × 10^−5^) (**Fig. 6C, D**). Significantly enriched iGAN genes, in contrast, were overrepresented in a neuron projection development network in both BIP and CUD (<u>Supplementary Figure 10</u> and <u>Supplementary Table 11</u> for q values of individual terms). The shared versus psychiatric disorder-specific pathway enrichments for iGLUTs are shown in <u>Supplementary Figure 11</u>, including a unique overrepresentation of iGLUT SEGs involved in “Synaptic Vesicle Cycle” in BIP (q = 7.25 × 10^−15^). Overall, we observed neuronal cell type-specific enrichment of risk variants for some psychiatric disorders, most notably CUD risk with iDANs and SCZ risk with iGANs and iGLUTs.

**Figure 6:**
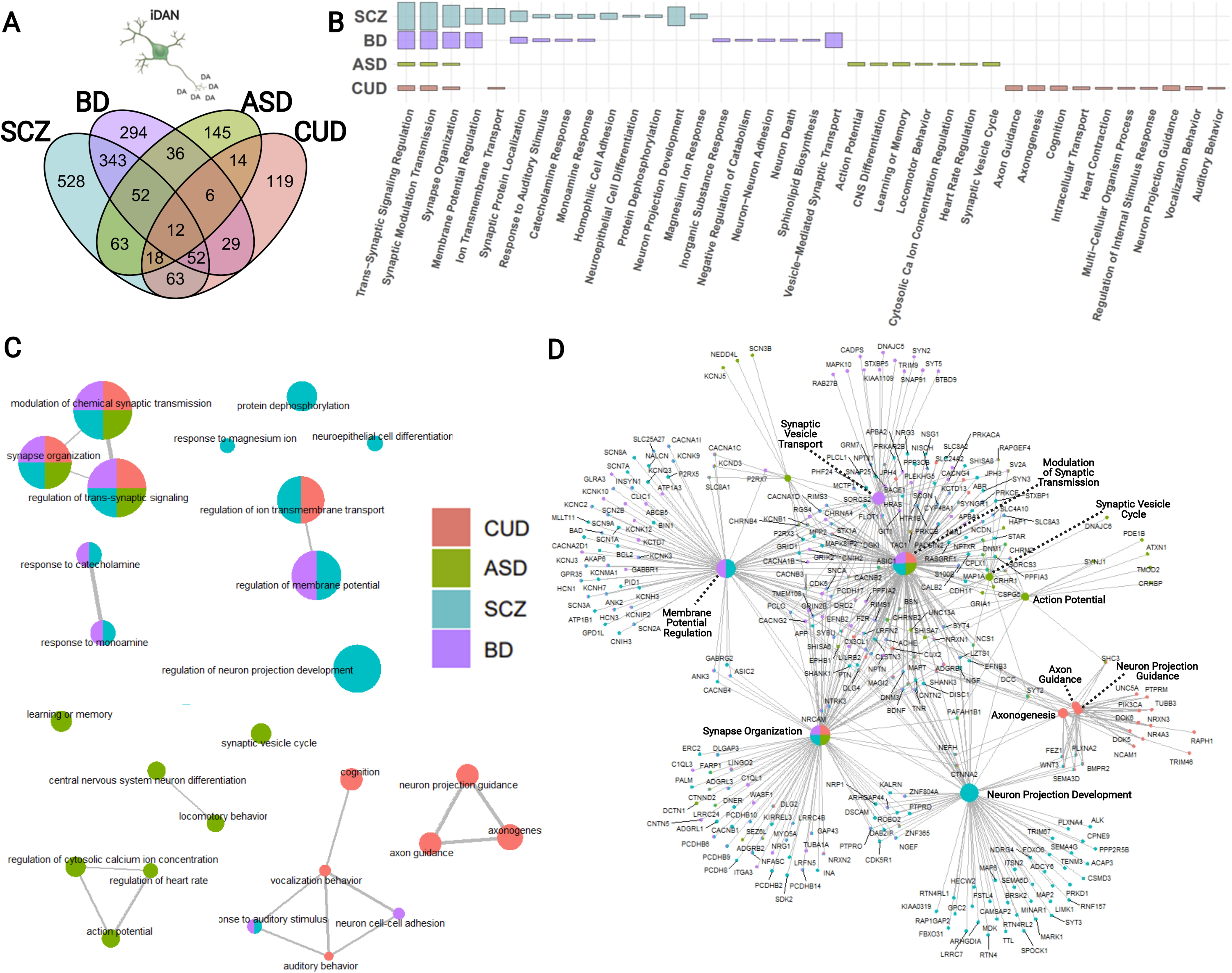
Biological interpretation of iDAN-specific gene expression with differential enrichment in genetic loci associated with schizophrenia (SCZ), bipolar disorder (BIP), autism spectrum disorder (ASD), and cannabis use disorder (CUD). (A) Venn diagram illustrating the overlap in significantly enriched iDAN SEGs among the four disorders. (B) Summary results from GSOA of enriched iDAN SEGs by disorder; height of the bar represents the magnitude of-log(FDR q values). (C) Enrichment map of top biological processes overrepresented among disorder-specific iDAN SEGs. Circle (node) colors indicate the disorder, and multiple colors within a node correspond to pathways shared among the indicated conditions. Node size represents the number of iDAN SEGs represented in the particular pathway. Pathway nodes are connected by edges to other nodes with shared genes to form pathway networks. The spatial arrangement between networks and free-standing nodes is not biologically meaningful and was adjusted for graphical purposes. (D) Gene-concept network plot depicting relationships between significantly enriched genes as a function of node membership among the top enriched pathways in iDANs. As in C, node size indicates the number of iDAN SEGs found within the GO set; colors of nodes and individual gene dots represent disorder involvement.

Finally, we investigated the extent to which correlation in genetic risk between psychiatric disorders was reflected in neuronal subtype-specific enrichment in psychiatric heritability. Broadly, disorder, rather than cell type, was the strongest driver of gene-level correlations in psychiatric heritability enrichments (<u>Supplementary Figure 14</u>; full results in <u>Supplementary Table 13</u>). Correlations between neuronal subtypes within each disorder were generally the lowest between iGANs and iGLUTs; for example, in CUD, the coefficient of correlation between iGANs and iGLUTs was *r* = 0.33 (p = 5.16 × 10^−152^), but *r =* 0.45 (p = 1.20 × 10^−306^) between iGANs and iDANs and *r =* 0.46 (p = 4.40 × 10^−313^) between iDANs and iGLUTs. A similar pattern was observed for SCZ and BIP (<u>Supplementary Figure 14</u>). For a given neuronal subtype, however, correlations between disorders were higher between SCZ and BIP than for any other disorder pair: for iDANs, the correlation between SCZ and BIP was *r =* 0.24 (p = 2.76 × 10^−81^) but only *r* = 0.070 (p = 5.97 × 10^−8^) between SCZ and ASD and *r* = 0.071 (p = 3.61 × 10^−8^) between SCZ and CUD, for example. These results suggest that the shared genetic risk architecture between SCZ and BIP^72–74^ is reflected in the heritability enrichments obtained from hiPSC-derived neuronal subtypes.

## DISCUSSION

Transient overexpression of just three transcription factors, *ASCL1, LMX1B,* and *NURR1*, yields homogenous populations of induced dopaminergic neurons. iDANs demonstrate a maturation-dependent increase in the expression of several marker genes of midbrain dopaminergic neuron identity and develop dopamine biosynthesis capabilities by DIV21. Relative to previous reports^10,11^, we demonstrate a substantial improvement in the yield and purity of iDANs. Moreover, we employed five independent donors to calculate percent purity, and note that the inter-donor variability (70-98%) was narrower than in a previous study that reported from 13-65% across four donors^12^. Our approach, which combined all three transcription factors and the selection cassette into a single doxycycline-inducible vector, ensured that the expression levels of all three transcription factors did not vary considerably relative to one another, as opposed to other methods where individual vectors deliver each transgene separately.

Although DIV35 iDANs are spontaneously active and exhibited hallmark physiological properties of *in vivo* MDNs, we did not detect I_h_ currents. These hyperpolarization-activated inwardly rectifying currents are mediated by HCN channel activity^75^. We found that *HCN1, HCN2, HCN3,* and *HCN4* were all expressed at high levels in iDANs (<u>Supplementary Figure 15</u>), and so it remains unclear why iDANs lack I_h_ currents. This likely reflects the immature nature of iDANs compared to adult MDNs, as I_h_ currents do not typically develop until the early post-natal period^76^ in rodents. iDAN cell capacitance, another measure of maturity, was 20 (±8) pF, comparable to values recorded in iGLUT neurons (22 ±1 pF)^77^ and human second trimester neurons (range of 18.5 ±2.5 pF to 24.8 ±3.5 pF)^78^. Likewise, competitive gene set testing also indicated global gene expression patterns consistent with an early neurodevelopmental, midbrain dopaminergic neuron phenotype. This is unsurprising, given previous reports that other types of hiPSCs-derived neurons most closely resemble human fetal neurons^79,80^, specifically those at 16-24 post-conception weeks^77^. This makes iDANs more suitable for studies of mechanisms related to psychiatric disease risk and onset, rather than phenotypes associated with late-stage disease.

Isogenic neuronal populations are uniquely suited for CRISPR-based functional genomic studies of subtype specific mechanisms across psychiatric disorders^81^. While CUD, BIP, and SCZ risk loci were enriched for unique subsets of specifically expressed genes in iDANs, iGANs, and iGLUTs, ASD was only enriched in iDAN SEGs. Genes enriched in psychiatric disease heritability in all three neuronal subtypes were overrepresented among biological pathways involved in synaptic structure and neurotransmission, implicating broad disruption of these processes in psychiatric disease. Different neuronal subtypes captured different aspects of disease biology; furthermore, within a given neuronal subtype, unique pathways were implicated in different disorders. Thus, we posit that each induced neuronal subtype captures both shared and distinct aspects of heritability enrichment that correspond to specific biological pathways that drive disease risk across different psychiatric conditions. Consistent with the shared genetic architecture of SCZ and BIP^72–74,82^, cross-disorder correlations were far greater between SCZ and BIP then between any other pair of conditions, although this effect may be somewhat inflated by overlapping control groups in SCZ and BIP GWA studies^51,56,83^. Moving forward, it will be critical to evaluate the functional effects of risk loci on gene expression and activity in specific subtypes of neurons in order to understand the mechanisms by which genetic variation adversely impacts brain phenotypes.

## FIGURE CAPTIONS

**<u>Supplementary Figure 1</u>:** RNAseq library-processing and quality control results using a standard pipeline^84^. (A) Raw sequencing data were mapped to a total of ~58,000 Ensembl gene IDs. Plotting the distributions of log_2_-transformed counts per million (CPM) values for all genes demonstrates that a substantial proportion of genes are expressed at low levels. (B) Removing lowly expressed genes produces a unimodal density plot of log_2_(CPM) values and enables downstream mean-variance relationships to be estimated with greater reliability^46^. Dashed line in (A) and (B) indicates the log_2_(CPM) cutoff value of −0.60, equivalent to about 0.66 CPM. (C) Boxplots showing the distributions of gene expression values across all RNAseq libraries prior to normalization. (D) Normalization using trimmed mean of M-values (TMM)^46^ adjusts libraries distributions using scaling factors that reflect differences in overall library sizes; this improves the similarity of expression distributions between libraries. (E) Mean-variance trend shows greater variability in lowly expressed genes in the filtered set of 16,641 genes. (F) Addition of precision weights produces a flat curve such that the variance is no longer dependent on the mean^20^.

**<u>Supplementary Figure 2</u>:** Longitudinal MEA analysis of iDANs shows maturation of physiological activity over the duration of the protocol. Local regression curves are fit with day in vitro (DIV) as the independent variable; the point estimates are drawn as a blue line with 95% confidence intervals surrounding in grey.

**<u>Supplementary Figure 3</u>:** Full results of competitive gene set testing for enrichment of hiPSC, iDAN, MDN, and non-MDN gene expression in specifically expressed genes for the 24 cell types in the K1 dataset^46^. Enrichment in the “down” direction is depicted by multiplying the - log(FDR q values) by −1.

**<u>Supplementary Figure 4</u>:** Full results for competitive gene set testing for enrichment of hiPSCs, iDANs, MDNs, and non-MDN gene expression in specifically expressed genes for the 149 cell types in the K2 dataset^46^. Enrichment in the “down” direction is depicted by multiplying the -log(FDR q values) by −1.

**<u>Supplementary Figure 5</u>:** Full results for competitive gene set testing for enrichment of hiPSC, iDAN, MDN, and non-MDN gene expression in specifically expressed genes for the 26 cell types from the La Manno et al (2016) dataset on the developing midbrain^20^. Enrichment in the “down” direction is depicted by multiplying the -log(FDR q values) by −1. DA, DA1, and DA2 = dopaminergic neuron subtypes; OMTN = oculomotor and trochlear nucleus neurons; RN = red nucleus neurons; SERT = serotonergic neuron; NMB = medial neuroblast; NBML1, NBML5 = mediolateral neuroblast types 1 and 5; GABA = GABAergic neurons; NBGABA = GABAergic neuroblast; NPROG = neural progenitor cells; PROGBP = basal plate progenitor cells; PROGFPL = lateral floor plate progenitor cells; PROGFPM = medial floor plate progenitor cells; PROGM = midline progenitor cells; RGL1, 2A, 2B, 2C, 3 = radial glia-like cells 1, 2A, 2B, 2C, and 3; BASAL = basal floor plate cells; ENDO = endothelial cells; MGL = microglia; OPC = oligodendrocyte precursor cells; PERIC = pericytes.

**<u>Supplementary Figure 6</u>**: Gene set testing results utilizing MAGMA-derived risk genes and post-mortem SEGs for MDNs and non-MDNs. Dashed line indicates nominal significance (p < 0.05), while * indicates those SEG sets that were significantly enriched after correction for multiple testing. Alzheimer disease = AD; attention-deficit/hyperactivity disorder = ADHD; anorexia nervosa = AN; autism spectrum disorder = ASD; alcohol use disorder = AUD; bipolar disorder = BIP; cannabis use disorder = CUD; cross-disorder pleiotropic loci = CxD; major depressive disorder = MDD; obsessive-compulsive disorder = OCD; opioid use disorder = OUD; Parkinson disease = PD; post-traumatic stress disorder = PTSD; schizophrenia = SCZ.

**<u>Supplementary Figure 7</u>:** Interpretation of iDAN, iGAN, and iGLUT SEGs significantly enriched in schizophrenia heritability. (A) Venn diagram showing overlap between significantly enriched genes among the three neuronal subtypes. (B) GSOA results implicating biological pathways overrepresented among neuronal subtype-enriched schizophrenia heritability. GO terms are organized to show pathways with shared enrichment among all three neuronal subtypes, followed by subtype-specific pathway enrichment in schizophrenia. (C) Gene-concept network plot depicting relationships among significantly enriched genes as a function of node membership among the top enriched pathways in iDANs, iGANs, and iGLUTs. Circle (node) colors indicate the neuronal subtype, and multiple colors within a node correspond to pathways shared among the indicated subtypes. Node size represents the number of SEGs represented in the particular pathway. (D) Enrichment map of shared and neuronal subtype-specific pathways enriched in schizophrenia heritability. Pathway nodes are connected by edges to other nodes with shared genes to form pathway networks. The spatial arrangement between networks and free-standing nodes is not biologically meaningful and was adjusted for graphical purposes.

**<u>Supplementary Figure 8</u>:** Biological interpretation of iDAN, iGAN, and iGLUT SEGs significantly enriched in bipolar disorder heritability. (A) Venn diagram showing overlap between significantly enriched genes among the three neuronal subtypes. (B) GSOA results implicating biological pathways overrepresented among neuronal subtype-enriched bipolar disorder heritability. GO terms are organized to show pathways with shared enrichment among all three neuronal subtypes, followed by subtype-specific pathway enrichment in bipolar disorder. (C) Gene-concept network plot depicting relationships of significantly enriched genes as a function of node membership among the top enriched pathways in iDANs, iGANs, and iGLUTs. Circle (node) colors indicate the neuronal subtype, and multiple colors within a node correspond to pathways shared among the indicated subtypes. Node size represents the number of SEGs represented in the particular pathway. (D) Enrichment map of shared and neuronal subtype-specific pathways enriched in bipolar disorder heritability. Pathway nodes are connected by edges to other nodes with shared genes to form pathway networks. The spatial arrangement between networks and free-standing nodes is not biologically meaningful and was adjusted for graphical purposes.

**<u>Supplementary Figure 9</u>:** Biological interpretation of iDAN, iGAN, and iGLUT genes significantly enriched in cannabis use disorder heritability. (A) Venn diagram showing overlap between significantly enriched genes among the three neuronal subtypes. (B) GSOA results implicating biological pathways overrepresented among neuronal subtype-enriched genes in cannabis use disorder heritability. GO terms are organized to show pathways with shared enrichment among all three neuronal subtypes, followed by subtype-specific pathway enrichment in cannabis use disorder. (C) Gene-concept network plot depicting relationships among significantly enriched genes as a function of node membership among the top enriched pathways in iDANs, iGANs, and iGLUTs. Circle (node) colors indicate the neuronal subtype, and multiple colors within a node correspond to pathways shared between the indicated subtypes. Node size represents the number of SEGs represented in the particular pathway. (D) Enrichment map of shared and neuronal subtype-specific pathways enriched in cannabis use disorder heritability. Pathway nodes are connected by edges to other nodes with shared genes to form pathway networks. The spatial arrangement between networks and free-standing nodes is not biologically meaningful and was adjusted for graphical purposes.

**<u>Supplementary Figure 10</u>:** Biological interpretation of iGAN specifically expressed genes with differential enrichment in genetic loci associated with schizophrenia (SCZ), bipolar disorder (BIP), and cannabis use disorder (CUD). (A) Venn diagram illustrating the overlap in significantly enriched iGAN SEGs among the three disorders. (B) Summary results from GSOA of enriched iGAN SEGs by disorder; height of the bar represents the -log(FDR q values). (C) Enrichment map of top biological processes overrepresented among disorder-specific iGAN SEGs. Circle (node) colors indicate the disorder, and multiple colors within a node correspond to pathways shared between the indicated conditions. Node size represents the number of iGAN SEGs represented in the particular pathway. Pathway nodes are connected by edges to other nodes with shared genes to form pathway networks. The spatial arrangement between networks and free-standing nodes is not biologically meaningful and was adjusted for graphical purposes. (D) Gene-concept network plot depicting relationships between significantly enriched genes as a function of node membership among the top enriched pathways in iGANs. As in C, node size indicates the number of iDAN SEGs found within the GO set; colors of nodes and individual gene dots represent disorder involvement.

**<u>Supplementary Figure 11</u>**: Biological interpretation of iGLUT specifically expressed genes with differential enrichment in genetic loci associated with schizophrenia (SCZ), bipolar disorder (BIP), cannabis use disorder (CUD), and cross disorder pleiotropic loci (CxD). (A) Venn diagram illustrating the overlap in significantly enriched iGLUT SEGs among the four disorders. (B) Summary results from GSOA of enriched iGLUT SEGs by disorder; height of the bar represents the -log(FDR q values). (C) Enrichment map of top biological processes overrepresented among disorder-specific iGLUT SEGs. Circle (node) colors indicate the disorder, and multiple colors within a node correspond to pathways shared between the indicated conditions. Node size represents the number of iGLUT SEGs represented in the particular pathway. Pathway nodes are connected by edges to other nodes with shared genes to form pathway networks. The spatial arrangement between networks and free-standing nodes is not biologically meaningful and was adjusted for graphical purposes. (D) Gene-concept network plot depicting relationships between significantly enriched genes as a function of node membership among the top enriched pathways in iGLUTs. As in C, node size indicates the number of iGLUTs SEGs found within the GO set; colors of nodes and individual gene dots represent disorder involvement.

**<u>Supplementary Figure 12</u>:** Biological interpretation of heritability enrichment of iDAN SEGs in autism spectrum disorder (ASD). (A) GSOA results implicating biological pathways overrepresented among iDAN SEGs enriched in ASD heritability. (B) Gene-concept network plot depicting relationships among iDAN SEGs as a function of node membership for the top enriched pathways. Node size represents the number of SEGs represented in the particular pathway. (C) Enrichment map of pathways implicated in iDAN specific gene expression in ASD enrichment. Pathway nodes are connected by edges to other nodes with shared genes to form pathway networks. The spatial arrangement between networks and free-standing nodes is not biologically meaningful and was adjusted for graphical purposes.

**<u>Supplementary Figure 13</u>:** Biological interpretation of heritability enrichment of iGLUT SEGs in cross-disorder pleiotropic loci (CxD). (A) GSOA results implicating biological pathways overrepresented among iGLUTs SEGs enriched in CxD heritability. (B) Gene-concept network plot depicting relationships among iGLUT SEGs as a function of node membership for the top enriched pathways. Node size represents the number of SEGs represented in the particular pathway. (C) Enrichment map of pathways implicated in iGLUT specific gene expression in CxD enrichment. Pathway nodes are connected by edges to other nodes with shared genes to form pathway networks. The spatial arrangement between networks and free-standing nodes is not biologically meaningful and was adjusted for graphical purposes.

**<u>Supplementary Figure 14</u>**: Pairwise correlations between MAGMA z-scores across all genes included in each set significantly enriched set. Color scale represents the magnitude of the correlation, and samples are ordered via hierarchical clustering.

**<u>Supplementary Figure 15</u>**: Gene expression values of *HCN1, HCN2, HCN3,* and *HCN4* in iDANs, MDNs, Non-MDNs, and hiPSCs showing highly similar expression levels of all four genes in iDANs and MDNs.

## Description of Supplementary Tables

<u>Supplementary Table 1</u>: Meta-data on both *in-vitro* and post-mortem donors and samples used for RNA-sequencing analyses

<u>Supplementary Table 2</u>: Sequences of forward and reverse primers used for qPCR assays

<u>Supplementary Table 3</u>: Product information and dilutions for primary and secondary antibodies used in immunocytochemical stainings

<u>Supplementary Table 4</u>: List of specifically expressed genes derived from the top 10% of gene *t*-statistics in hiPSCs, iDANs, iGANs, and iGLUTs

<u>Supplementary Table 5:</u> Specifically expressed genes from the 24 K1 cell types^46^, the 149 K2 cell types^46^, and the 26 developmental midbrain cell types^20^ used for competitive gene set testing

<u>Supplementary Table 6</u>: GWAS datasets used for MAGMA analyses of neuronal subtype-specific heritability enrichment

<u>Supplementary Table 7</u>: Summary statistics for results of electrophysiology recordings

<u>Supplementary Table 8</u>: Results of gene-set overrepresentation analyses of differentially expressed genes in hiPSCs, iDANs, MDNs, and non-MDNs among KEGG pathways and GO Biological Process terms

<u>Supplementary Table 9</u>: Results of competitive gene set testing of differentially expressed genes in hiPSCs, iDANs, MDNs, and non-MDNs among the K1^46^, K2^46^, and La Manno^20^ datasets

<u>Supplementary Table 10</u>: Results of MAGMA-based heritability enrichment analyses, including the summary statistics of each geneset test as well as the significantly enriched genes for each disorder with one or more heritability enrichment in a neuronal subtype

<u>Supplementary Table 11</u>: Complete results of gene-set overrepresentation analyses of significantly enriched genes from MAGMA testing of heritability enrichments in neuronal subtype-specific gene expression

<u>Supplementary Table 12</u>: Summary statistics for all genes in significantly enriched sets from MAGMA-based analyses

<u>Supplementary Table 13</u>: Results of correlation analyses

## Acknowledgements

This research was supported by R01MH106056, U01DA047880, R01DA048279, 6R56MH101454. Figures in this manuscript were created with Biorender.com. The authors wish to thank Rachel Oren for helpful feedback on an earlier version of this manuscript and Dr. Stefano Marenco, Dr. Barbara Lipska and Dr. Pavan Auluck and their staff in the Human Brain Collection Core at the National Institutes of Health for providing postmortem brain tissues.

## Author Contributions

SKP, SA, and KJB conceived of the study. SKP, KT, IP, KD, PS, LMH, SA, and KJP designed experiments. SKP, COS, IP, KD, RE, and SH conduced experiments. MI, TL, and AV performed FANS of post-mortem samples. SKP, COS, MI, TL, and AV prepared RNA-sequencing libraries. SKP, KT, and WL conducted computational and bioinformatic analyses. SKP wrote the paper, with contributions from KT and KD. All authors reviewed the manuscript and approved of it in its final form.

## Supplementary Note 1

*Detailed Protocol for Production of iDANs*

Unless otherwise noted, the concentration of chemicals added to media will remain identical to that seen at the first time in which it was mentioned. For example, when the author writes “doxycycline 1 ug/ml” on “step a,” and just “doxycycline” in subsequent steps, it should be assumed that the concentration of doxycycline is still 1ug/ml unless specified otherwise.

## Protocol for Induced Dopaminergic Neurons (iDANs)

- DIV0: harvest hiPSCs in Accutase to obtain a single-cell suspension of iPSCs. Quench Accutase suspension with DMEM at a volumetric ratio of 1:3 (Accutase:DMEM). Spin at RT for 5 minutes at 800g. Once pelleted, aspirate the supernatant and resuspend the pellet in 1.0 mL of StemFlex with Y27632 ROCK inhibitor (StemCell Technologies, #72302). Count the cells. Then, dilute using StemFlex with ROCK Inhibitor to a concentration of 1e6 (1 million) cells per mL and a final ROCK inhibitor concentration of 10uM. Then, add the appropriate viruses, *rTTa* and *ALN-PuroR*. See trouble-shooting recommendations for further notes on amount of virus to add.

- <u>Technical note</u>: Ensure that your viruses are at an appropriate concentration of 1e7 IU/mL using a qPCR Lentivirus Titration Kit (Applied Biological Materials, #LV900)
- Plate the StemFlex-ROCK Inhibitor-hiPSCs-virus suspension on matrigel-coated (40ug/mL, but see below) six-well plates at a density of 1.5e6 cells per well of a six well plate. Incubate overnight (at least 12 hours, preferably 16-24, but never greater than 48).

- Trick: We have found that the cells appear healthier and that there is a dramatic reduction in “flat cells” by starting from DIV0 with plating the cells on plates coated with 160ug/mL of matrigel.
- <u>Technical note</u>: for each well of a 12 well plate, seed 500k-750k per well; for a 24 well, seed 250k per well.
- <u>Tip</u>: If, after your first batch with a given cell line, you find that it forms many flat cells, one thing to try is to plate only 1 million cells per well of a six well plate. In this case, add at least 0.5mL of additional StemFlex with THX Rock Inhibitor
- After the overnight incubation, aspirate the media and replace with <u>Induction Media</u> (see recipe below) with doxycycline 1ug/mL. This is **<u>DIV1.</u>**
- The next day, on **<u>DIV2</u>**, replace the media with Induction Media with doxycycline 1ug/mL and Puromycin 2ug/mL.
- The next day, on **<u>DIV3</u>**, repeat the prior step, as there are often many dead cells. The next day, on **<u>DIV4</u>**, you may either (a) keep the media as is, (b) change to the same media as in the previous step if there are many flat cells, or (c) if, and only if, there are processes beginning to develop across most cells, replace the media with induction Media with Doxycycline, puromycin, and 2μM Ara-C.

- It can be challenging to select a precise, ideal time to add Ara-C. This may vary from cell line to cell line and across different batches of virus. A balance must be struck between making every effort to prevent the development of flat cells and not killing your “good” cells that eventually become iDANs. As a general rule of thumb, once you see that most of the cells on the plate have early processes that have begun to extend from the cell body, you should start Ara-C.
- **<u>DIV5:</u>** On this day, you may do the same thing as written in the previous step
- **<u>DIV6</u>**: At this point, the cells have undergone 4 days of puromycin selection. That is enough. Replace the media such that it does not contain puromycin (still <u>Induction Media</u>). It is also at this time that Ara-C 2μM should be initiated at the latest. If the cells are a solid sheet of flat cells with <10% of cells with the desired morphology, then the experiment is not likely going to work. Go to the troubleshooting section and try again.
- **<u>Guidelines for splitting cells:</u>** There is no definitive, exact day on which to split the cells and replate. See the recommendations that shortly follow below. Upon splitting and replating, aspirate the media and incubate in 1.0mL Accutase per well at 37 degrees for at least 5 minutes but no greater than 20 minutes; quench with DMEM at a volumetric ratio of 1:3 Accutase:DMEM. Spin down at 1000g for 5 minutes at room temperature. Aspirate the supernatant, resuspend in 1 mL induction media with ROCK inhibitor, doxycycline, and Ara-C. Count. Dilute to a concentration of 1e6 cells/mL. Replate onto plates double-coated with 0.1% PEI (first) + 80ug/mL matrigel OR 160ug/mL matrigel only.

- <u>Tip</u>: the decision about when to split and replate should be guided by a few important factors. (1) splitting and plating on PEI helps to kill off flat cells to some extent, although it will not completely solve the problem; (2) you do not want to split so early that your final plate (PEI and matrigel coated) ends up having flat cells develop because you have not adequately killed them off and (3) you do not want to split so late that the cells are more mature neurons and they die due to the stress of splitting. It is generally helpful for the cells to have had at least 24 hours, preferably 48, of Ara-C exposure. Doing so will have killed or weakened the flat cells such that when you do split, they die off and do not re-plate.
- Do not replate past DIV14. That is the latest you should replate.

- **on average, across cell lines and batches, the DIV on which is split ranges from DIV5-9.**
- <u>Technical note</u>: make sure to have prepared your plates prior to splitting. To coat PEI/matrigel: add 2.0mL of 0.1% PEI in borate buffer solution to each well of a 6 well plate. Incubate at 37 C for exactly 60 minutes. Then, aspirate PEI and wash 5 times with pure water. Do not wash fewer than 5 times. PEI is toxic and residual PEI may lead to excessive cell death. Then, add 2mL of 80ug/mL matrigel in each well. Incubate for at least 30 minutes.
- <u>Technical note</u>: upon re-platting, seed at a density of 1-3 million cells per well of a six-well plate. If there were significant levels of flat cells when you split at the Accutase step, choose the lower end of 1e6 cells per well. If there were none, you can plate 3 million (no greater than 5e6 per well).
- **<u>Split the cells</u>:** Using the criteria mentioned above, make sure to have split the now iDAN progenitors by DIV14 at the absolute latest.
- Keep in Induction Media and Doxycycline until DIV14
  - Keep Ara-C in the media up until you no longer see flat cells and then keep it in for two additional days at a concentration of at least 1μM. However, do not keep Ara-C in the media for greater than 10 days. That starts to make the good, future dopaminergic neurons unhealthy.
- At **<u>DIV14:</u>** switch to <u>Neuron Media</u> without doxycycline

- Perform half media changes every other day until the desired time point

## MEDIA RECIPES

**Induction Media**: 500 mL DMEM F12 with Glutamax and Sodium Pyruvate (already in media); 5 mL Anti-Anti; 5 mL N2; 10mL B-27 without vitamin A; 500 uL doxycycline for a final concentration of 1ug/mL

**Neuron Media:** BrainPhys; 1:100 Anti-Anti, Glutamax, Sodium Pyruvate, N-2; 1:50 B-27 without vitamin A; 20ng/uL BDNF, 20ng/uL GDNF, 200nM Ascorbic Acid, 500ug/mL dibutyryl cAMP; 1ug/mL mouse laminin

### Trouble-Shooting the Emergence of Too Many “Flat Cells”

Flat cells are cells that survive antibiotic selection but do not become neurons. These are either dividing cells or apoptotic cells generated from being too harsh in your treatment of the cells during selection or due to too much virus. The following may be attempted if you run into issues with flat cells:

1. Double the puromycin dose (including on DIV2). This raises the threshold required to survive.
2. Use 4μM Ara-C instead of 2μM.
3. If flat cells are showing up later on in your differentiation, keep Ara-C in longer (but not to exceed 10 days)
4. When plating the hiPSCs with virus on DIV0, decrease the density of cells. From anecdotal experience, when cells are very confluent, they do not differentiate well
5. Use 160ug/mL matrigel throughout the entire experiment, starting at DIV0.
6. Decrease the amount of ALN-PuroR virus (start by a 50% reduction in the volume added, assuming you’re using a consistent batch of virus). This is a highly effective way to decrease the overall burden of any multi-protein transgene products produced due to incomplete cleavage between 2a peptide sequences. Note, this may increase the number of dead cells observed with puromycin selection.

### Reagents

- BrainPhys: StemCell Technologies, #05790
- DMEM F12+ with Glutamax and Sodium Pyruvate: Thermo Fisher, #10565018
- Matrigel, growth factor reduced: Corning #354230
- Polyethylenimine: Sigma/Millipore #P3143
- Pierce Borate buffer 20X (dilute 1:10, then make 0.1% PEI solution): Thermo Fisher #28314
- DMEM: Thermo Fisher #11966025
- BDNF: Peprotech, #45002 (media concentration of 20 ng/mL)
- GDNF: Peprotech, #45010 (media concentration of 20 ng/mL)
- cAMP: Sigma #D0627 (media concentration of 500 ug/mL)
- Ascorbic Acid: Sigma #A0278 (media concentration of 200nM)
- N2: Thermo Fisher, #17502-048
- B27, Thermo Fisher
- Natural mouse laminin, Thermo Fisher #23017-015 (media concentration of 1ug/mL
- Rock inhibitor Y-26732 (media concentration of 10uM)
- StemFlex, Thermo Fisher #A3349401
- Accutase: Innovative Cell Technologies #AT104
- Antibiotic-antimycotic: Thermo Fisher #15240062
- Glutamax: Thermo Fisher #35050061
- Sodium pyruvate: Thermo Fisher #11360070

